# High extraversion enhances attentional control through dynamic network reorganization

**DOI:** 10.1101/2025.11.20.689570

**Authors:** Joseph CC Chen, Bijurika Nandi, Richard Campusano, Fei Jiang, David Ziegler, Adam Gazzaley, Theodore Zanto

**Affiliations:** Neuroscape, Department of Neurology, University of California San Francisco, CA 94158; Department of Epidemiology and Biostatistics, University of California San Francisco, CA 94158; Departments of Physiology and Psychiatry, University of California San Francisco, CA 94158

**Keywords:** Extraversion, fMRI, dynamic functional connectivity, attention, meditation

## Abstract

The extraversion-introversion dimension of personality is hypothesized to differ based on low or high cortical arousal, respectively. Notably, high cortical arousal in introverts is thought to underlie increased distractibility. Here, we assess fMRI while participants meditate (focused attention to their breath) under three levels of auditory distraction: no, low and high. Whereas introverts exhibited worsening attentional focus on their breath with increasing distraction, extraverts retained their ability to focus attention despite distraction. Dynamic functional connectivity analysis indicated that extraverts exhibited less globally efficient and less modular networks, which may prevent distracting stimuli from creating interference. Furthermore, connectivity strengths amongst the default mode, central executive, and salience networks were increased for extraverts and decreased in introverts during high focused attention; potentially indicating distinct cognitive processes that support attentional control. These findings support the hypothesis regarding differing levels of cortical arousal in extraverts and introverts and extend personality theory by linking the extraversion dimension to attentional control and functional connectivity dynamics.

## 1. Introduction

Extraversion and introversion refer to personality types that categorize individuals as those who orient more towards the external or internal world, respectively. As such, people who are more extraverted are thought to be more social and outgoing, whereas those who are more introverted are more comfortable focusing on their own thoughts and ideas. The extraversion-introversion dimension is included in many theories of personality such as the five-factor model (McCrae & John, 1992) and Eysenck’s three-factor model (Eysenck & Eysenck, 1969). Extraverts are thought to also exhibit lower cortical arousal (i.e., global levels of cortical excitation/inhibition) compared to introverts (Eysenck, 1963). As it has been proposed that all individuals aim to achieve moderate levels of cortical arousal, extraverts may seek external sensations to achieve an optimal level of cortical arousal, whereas introverts avoid external sensations to achieve optimal levels (Eysenck & Eysenck, 1969).

Introverts’ higher cortical arousal levels have been interpreted as higher cortical excitability, which translates to lower sensory thresholds – demonstrated by the fact that introverts have lower auditory thresholds (Smith, 1968), lower visual sensitivity (Siddle et al., 1969), and lower somatosensory thresholds (via electrocutaneous stimulation) (Edman et al., 1979). Further investigations have linked these lower somatosensory thresholds to increased neural activity in the primary somatosensory cortex (Schaefer et al., 2012). These personality differences in cortical arousal and sensory thresholds may translate to differences in cognition – specifically in attentional control – due to the differing sensitivities to sensory stimuli.

Extraverts typically exhibit greater levels of attentional control (Hahn et al., 2015) that underlies the ability to control or direct their attention is the ability to resist distraction or competing information – often termed inhibitory control (Munakata et al., 2011). Using Stroop tasks to assess inhibitory control, extraverts demonstrate increased ability to ignore task-irrelevant information (Prabhakaran et al., 2011). When including distracting background high arousal music or noise, extraverts also performed better than introverts on the Stroop task (Cassidy & MacDonald, 2007). In another study, extraverts and introverts participated in comprehension and problem-solving tests in silence or in the presence of music. Extraverts performed better with music and worse in silence, whereas introverts performed better in silence and worse with music (Mistry, 2015). In the presence of background white noise, extraverts’ selective attention was improved compared to introverts (Moradi et al., 2019). The selective attention task used tested whether participants could detect a certain shape (downward-pointing triangle) was present amongst seven presented triangles (Moradi et al., 2019). Taken together, these results suggest that extraverts are more capable than introverts in resisting the negative impacts of external distractors.

While research supports the idea that introverts have more difficulty ignoring environmental distractions, it remains unclear whether this finding is attributable to differences in cortical arousal. As modern neuroimaging methods have investigated the links with cortical arousal and cognition, we attempt to bridge the knowledge gap between psychological hypotheses of personality traits, cognitive science of inhibitory control, and the neuroimaging of brain networks.

The functional magnetic resonance imaging (fMRI) blood oxgygenation-level-dependent (BOLD) signal has been used to track levels of cortical arousal (Chang et al., 2016; Liu et al., 2018) and underlying large-scale brain network patterns (Young et al., 2017). For example, cortical arousal via the ascending arousal system is functionally connected to the default mode network (DMN) and salience network (SN), and these connections are mediated by other traits such as age (Guardia et al., 2022). Beyond connectivity analyses, graph theoretical analyses of large-scale brain networks have further delineated between high and low arousal states with low arousal states associated with reduced network modularity and increased global connectivity (Lee et al., 2022). This relationship suggests that fMRI connectivity and graph analyses may be able to index cortical arousal, and by extension, personality trait differences. Indeed, prior work linking extraversion to resting-state fMRI (rsfMRI) and connectivity analyses have identified the involvement of the DMN and SN. For example, rsfMRI connectivity within the DMN (including precuneus and bilateral superior parietal lobes) were associated with extraversion (Sampaio et al., 2014), while strong engagement of the precuneus was associated with introversion (i.e., low extraversion) (Mulders et al., 2018). As such, DMN connectivity may correspond to Eysenck’s proposal of cortical arousal, whereby extraverts purported lower arousal corresponds to lower DMN connectivity, and this may be associated with decreased distractibility Conversely, increased SN recruitment is associated with higher extraversion (Tian et al., 2018), potentially reflecting increased external distractibility.

Together, previous research has shown that both functional connectivity and inhibitory control are differentially affected by personality traits. Here we explore whether there is a relationship between differential functional connectivity states and inhibitory control abilities. To elucidate these mechanisms, we use the focused attention on breath (meditation) paradigm, as the neural mechanisms associated with this task have been well-characterized.

Focused attention meditation (e.g., sustained focusing on one’s breath) engages attentional resources. During focus-on-breath meditation, the default mode network (DMN) is deactivated (e.g., the posterior cingulate cortex) (K. A. Garrison et al., 2013; Hasenkamp & Barsalou, 2012), and the central executive network (CEN) is activated (Hasenkamp & Barsalou, 2012). Furthermore, the salience network (SN) is associated with awareness of mind wandering (Hasenkamp & Barsalou, 2012). The interplay between these three key networks underlies the cognitive processes elicited by the focus-on-breath paradigm. Indeed, the interplay between the triple networks’ fMRI signals (DMN, CEN, and SN) appear to be real-time indicators of the focus-on-breath paradigm (Kim et al., 2019). Importantly, the focus-on-breath paradigm recruits brain areas previously implicated in extraversion and presents a promising paradigm to further understand the mechanism by which extraverts’ and introverts’ inhibitory control differ.

The present study used a focus-on-breath meditation paradigm while music was (or was not) present in the background. To further manipulate the level of environmental distraction, two types of music were selected: low and high arousal. While engaged in this focus-on-breath meditation task, fMRI data was collected. Participants were determined to be extraverts or introverts according to self-reported responses on the big five inventory personality questionnaire (McCrae & Costa Jr., 1999). In line with prior research, we hypothesized that when compared to introverts, extraverts would exhibit superior inhibitory control as revealed by a better ability to focus on their breath despite the presence of background music. To discover the neural differences possibly underlying these hypothesized cognitive differences, we assessed how functional connectivity within the DMN, CEN, and SN might differ between personality types. To delineate the various dynamic states that occur over time, we employed a dynamic functional connectivity analyses technique alongside graph theoretical analysis to characterize the effects of differing cortical arousal states which may uncover underlying neurophysiological differences between extraverts and introverts.

## 2. Methods

Participants gave informed consent in accordance with the Institutional Review Board of the **REMOVED_FOR_MASKED_REVIEW** and received a small fee in compensation for their time. This study was not preregistered. The analysis code is publicly available at: **REMOVED_FOR_MASKED_REVIEW.**

### 2.1. Participants

Thirty-seven participants underwent MRI sessions. All participants had normal or corrected-to-normal vision and normal hearing. One participant was excluded due to incomplete data collection from the scan. Two participants were excluded due to task noncompliance. Participants completed the big five inventory personality questionnaire based on the five factor model of personality (McCrae & Costa Jr., 1999) – which contains 8 questions (answerable on a 5-point Likert scale) dedicated to the introversion-extraversion scale which are shown in Table S1. Participants who scored: <24 were classified as introverts (N=17), >24 were extraverts (N=16), and =24 were excluded (N=1) leaving a total of 33 participants who completed the full fMRI paradigm and were included in the analysis.

**Table 1.** Participant details and demographics. Continuous variables are summarized as mean ± standard deviation.

| Demographic | Introverts | Extraverts |
| --- | --- | --- |
| Number | 17 | 16 |
| Age | 27.4 ± 5.7 | 26.7 ± 5.0 |
| Age Range | 20 – 38 | 21 – 36 |
| Sex (Male / Female) | 5 M, 12 F | 4 M, 12 F |
| Handedness (Left / Right) | 2 L, 15 R | 1 L, 17 R |
| Five Factor Model of Personality Scores (5-point Likert Scales, 1 – 5) |  |  |
| Extraversion Score (8 – 40) | 20.2 ± 2.3 | 30.9 ± 3.8 |
| Agreeableness Score (9 – 45) | 33.8 ± 5.1 | 37.4 ± 4.2 |
| Conscientiousness Score (9 – 45) | 34.5 ± 5.3 | 34.4 ± 5.0 |
| Neuroticism Score (8 – 40) | 25.6 ± 5.6 | 23.9 ± 6.7 |
| Openness Score (9 – 45) | 36.9 ± 4.0 | 39.9 ± 5.7 |

### 2.2. Paradigm

Each participant was instructed to listen to 169 25-second song clips at home and asked to rate each clip with valence and arousal on a visual analogue scale. Twelve songs rated by participants as high arousal (HA) and twelve rated as low arousal (LA) were selected for the following MRI paradigm.

Seven MRI scans per participant were collected at the **REMOVED_FOR_MASKED_REVIEW** on a 3T Prisma Fit: one high resolution anatomical scan and six fMRI (task-based) scans. The anatomical scan was T1-MPRAGE with the following parameters: TR = 2300 ms, TE = 2.9 ms, TI = 900 ms, flip angle = 9°, 1 mm isotropic voxels (160 x 240 field of view), 256 z-slices; and six multiband fMRI scans: TR = 850ms, TE = 3.28ms, multiband factor = 6, phase acceleration factor = 3, flip angle = 45°, 2.2 mm isotropic, 450 volumes collected, 96 x 96 x 66 dimensions collected, with 10 fMRI volumes discarded prior to the commencement of the 450 volumes (382.5 seconds). The six fMRI scans were randomized orders of AudioHA, AudioLA, BreathHA, BreathLA, MindWander, and Meditation scans. In each fMRI scan, participants were presented with twelve 30-second trials of: 25 seconds of “task” and 5 seconds to “respond” – with the “task” differing depending on each scan type. In the 5s response section, a question (depending on scan type) was presented with a visual analogue scale with values ranging from “Never” (left position) to “Always” (right position). A pointer, which always started at the middle (“Half the time”), was movable using a button box (Current Designs, PA) with two buttons moving the pointer towards the left or right respectively.

The “Audio” task (for both AudioHA and AudioLA) instructed participants to focus on the music clips (which were 25 seconds long each). The twelve trials in each scan (AudioHA and AudioLA) corresponded to the twelve songs previously rated by each participant for both HA and LA. The response section posed the question: “How much were you focused on the audio?”.

The “Breath” task (for both BreathHA and BreathLA) instructed participants to focus on their breath while the same twelve songs in the corresponding Audio task was played to participants. The response section posed the question “How much were you focused on your breath?”.

The “Meditation” task instructed participants to focus on their breath while no music played in the background. The response section similarly posed the question “How much were you focused on your breath?”.

The “MindWander” task instructed participants to let their mind wander while no music played in the background. The response section posed the question “How often were you mind wandering?”.

The present study only reports the results from the BreathHA, BreathLA, and Meditation scans due to the same response question being asked. This allows a direct comparison for how environmental distractions (i.e., background music) may affect attentional control.

### 2.3. fMRI Preprocessing and Analysis

All MRI scans were organized into Brain Imaging Data Structure format, preprocessed using fMRIprep v23.1.3 (Esteban et al., 2019), and postprocessed with XCP-D v0.6.0 (Mehta et al., 2024) using 24 motion estimates (“24P”). Dynamic functional connectivity was performed using time-varying dynamic network (TVDN) modelling (Jiang et al., 2022) which identifies dynamic state transitions within each fMRI scan. The Schaeffer 100 atlas with 56 subcortical regions parcellation output from XCPD was used for TVDN analysis. To perform successful TVDN modelling, parameters *r* (from 0.5 to 0.9 in 0.1 increments) and *kappa* (from 1.0 to 4.0 in 0.1 increments) were tuned to minimize the residual mean square error across all scans and all participants. The final parameters which yielded the lowest error were *r* = 0.5 and *kappa* = 1.6 which identified the number of state switches per scan.

The functional connectivity matrices between each state transition were further obtained as previously described (Jin et al., 2023). Briefly, the TVDN method models the presence of multiple (typically, 6 – 10) resting-state networks in each scan which dynamically occurred with different weightings across time. Functional connectivity matrices were generated by calculating the relative weightings (eigenvalues) of each underlying resting-state network. Each undirected (symmetrical) functional connectivity matrix (which contains z-transformed edge strengths) was then loaded into Matlab v2023a (MathWorks, MA), thresholded and binarized at the top 20% of connections, and analyzed for graph metrics using the Brain Connectivity Toolbox (Rubinov & Sporns, 2010) – specifically global efficiency (*efficiency_bin.m*) and modularity (*modularity_und.m*). Each fMRI scan would on average yield 21.4 state transitions – and subsequently, ∼22 matrices and ∼22 measures of global efficiency and modularity per scan. For each fMRI scan, the values of global efficiency and modularity were averaged to yield one value of global efficiency and modularity per scan. This would give an indication of the types of network organizations that each individual’s brain would tend towards in the given tasks. Finally, linear mixed effect models were used to predict the mean attention (across twelve trials) using fixed effects of graph metrics and Introvert/Extravert factor – including interaction, with each individual modelled as a random effect: *Mean Attention ∼ Graph Metric * Extravert + (1|ID)*, with graph metric either being a global efficiency or modularity. The interaction effect was assessed by a type 3 analysis of variance via “car” package (Fox & Weisberg, 2019) and the subsequent intercept and differing slope coefficients graphically presented via “ggplot2” (Wickham, 2016). Where relevant, the *post hoc* tests were performed using the “glht” from the “multcomp” package (Hothorn et al., 2008). However, for ease of interpretation, all introverts’ and all extraverts’ mean attention and graph metrics were Pearson correlated.

While changes in global efficiency and modularity were observed, it was unclear which functional network was responsible for these global changes. As such, two further analyses were performed. Firstly, the average number of top 20% edge strengths was classified as internal (i.e., edges connecting nodes of the same network) or external (i.e., edges connecting nodes of different networks). Furthermore, the number of inter-network edges connecting between the CEN, DMN, and SN were also investigated. The number of these internal and external edges pertaining to the CEN, DMN, and SN were calculated per scan and submitted to linear mixed effect modeling. Secondly, the strength of the top 20% edge strengths were analyzed. The average of the Fisher’s z-transformed values of the top 20% edge strengths for each network were analyzed for internal and external edges pertaining to the CEN, DMN, and SN and submitted to linear mixed effect modeling. For both the number of top 20% edges and the edge strengths, the interaction effect with extraversion personality was of interest to observe whether personality influenced network connectivity.

## 3. Results

### 3.1. Introverts exhibit reduced inhibitory control

To assess whether distraction affected participants’ ability to focus on their breath, self-reported measures of attentional focus were submitted to a linear mixed effect model with random intercepts and false discovery rate (FDR)-corrected *post hoc* tests. Participants rated their attentional focus in response to the question “How much were you focused on your breath?” on visual analogue scales ranging from “Never” (left position, value = 0) to “Always”(right position, value = 1). Main effects were observed from background music type (χ*^2^* = 29.5, *p* = 3.9e-7) and personality type (χ*^2^* = 4.5, *p* = 0.033), revealing that the no music condition and extraverts corresponded with the highest focus on breath. Furthermore, a significant interaction between background music type and personality type was observed (χ*^2^* = 7.6, *p* = 0.023), such that extraverts focused on breath significantly better in the presence of high arousal (HA) music compared to introverts (*t* = 2.13, *df = 35.9*, *p* = 0.032; Figure 1). Introverts reported a step-wise increase in their ability to focus on breath based on the level of distraction: low arousal (LA) music was more distracting than no music (*t* = 2.59, *df = 1151*, *p* = 0.0098; Figure 1), and HA music was more distracting than LA music (*t* = 2.84, *df = 1151*, *p* = 0.0045). Comparatively, extraverts reported higher focus in the no music condition compared to the LA music condition (*z* = 2.14, *p* = 0.033), but no difference between HA or LA music (*z* = 0.36, *p* = 0.72). Together, these results suggest that introverts were more negatively affected by environmental distractions while meditating.

**Figure 1.**
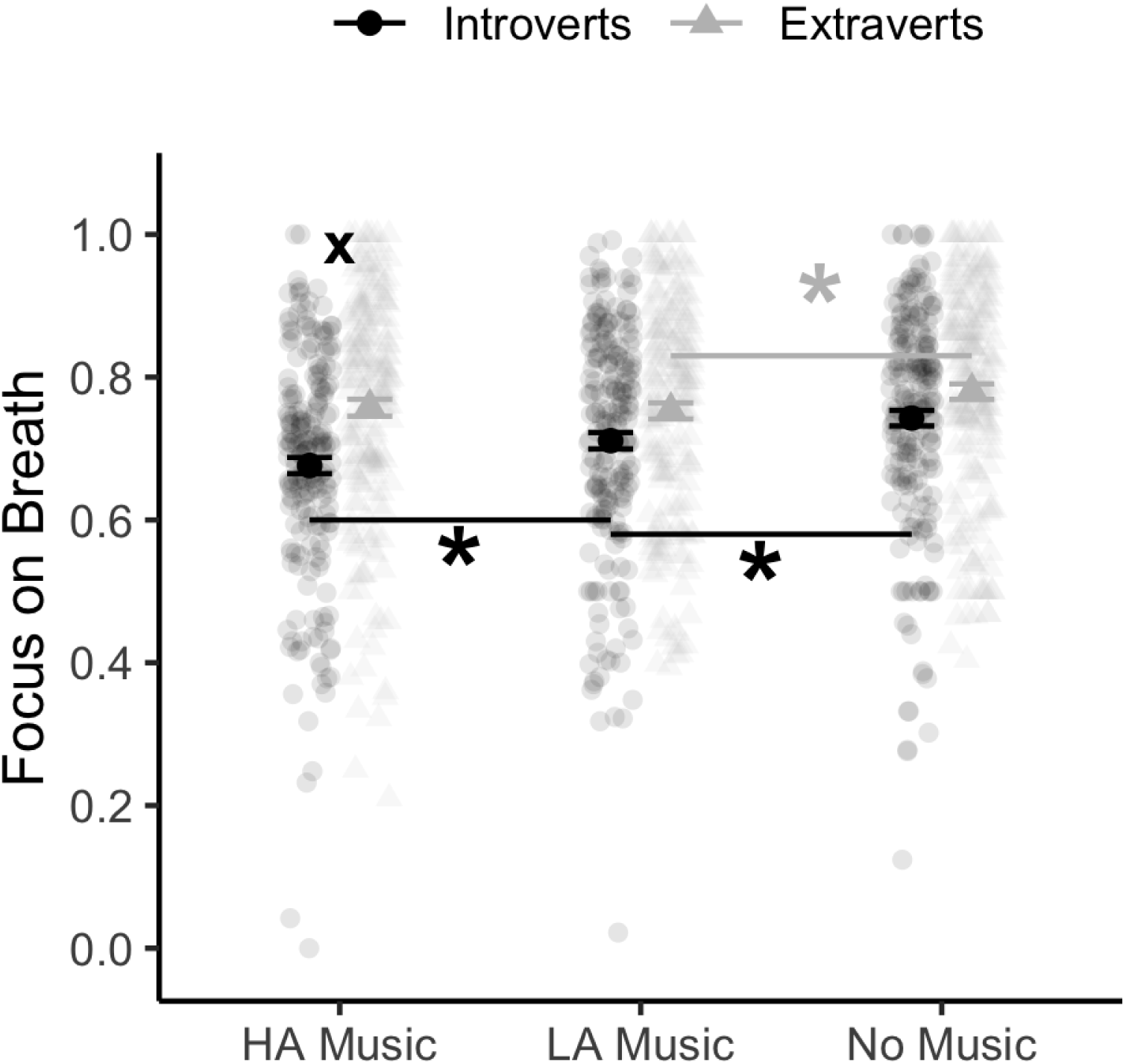
Attention Scores during Focus-on-Breath Task while No Music, Low Arousal music (LA), and High Arousal music (HA) is played to introverted (black, circle) or extraverted (grey, triangle) individuals. Solid points denote mean and error bars denote standard error. Semi-transparent points indicate individual trial data. Asterisks (*) and crosses (x) denote significant (p<0.05) false discovery rate corrected differences within-group and between-group respectively.

### 3.2. Extraverts reorganize networks to sustain attention

Graph theoretical analysis of the dynamic functional networks observed in participants while performing the paradigm also showed differences between introverts and extraverts. Specifically, linear mixed effect modeling showed interaction effects in global efficiency (χ *^2^* = 7.61, *p* = 0.0058, Figure 2A) and modularity (χ*^2^* = 4.32, *p* = 0.038, Figure 2B) between introverts and extraverts. As extraverts reported increasing ability to focus on their breaths, the functional networks were lower in global efficiency and modularity (Figure 2). Conversely, introverts exhibited opposite trends whereby increasing focus on breath were associated with higher global efficiency and higher modularity (Figure 2). However, if introverts were separately analyzed via a Pearson correlation, these effects show small correlation coefficients (Figure 2). Therefore, this interaction was largely driven by the extraverts’ relationship between attention and dynamic functional network metrics.

**Figure 2.**
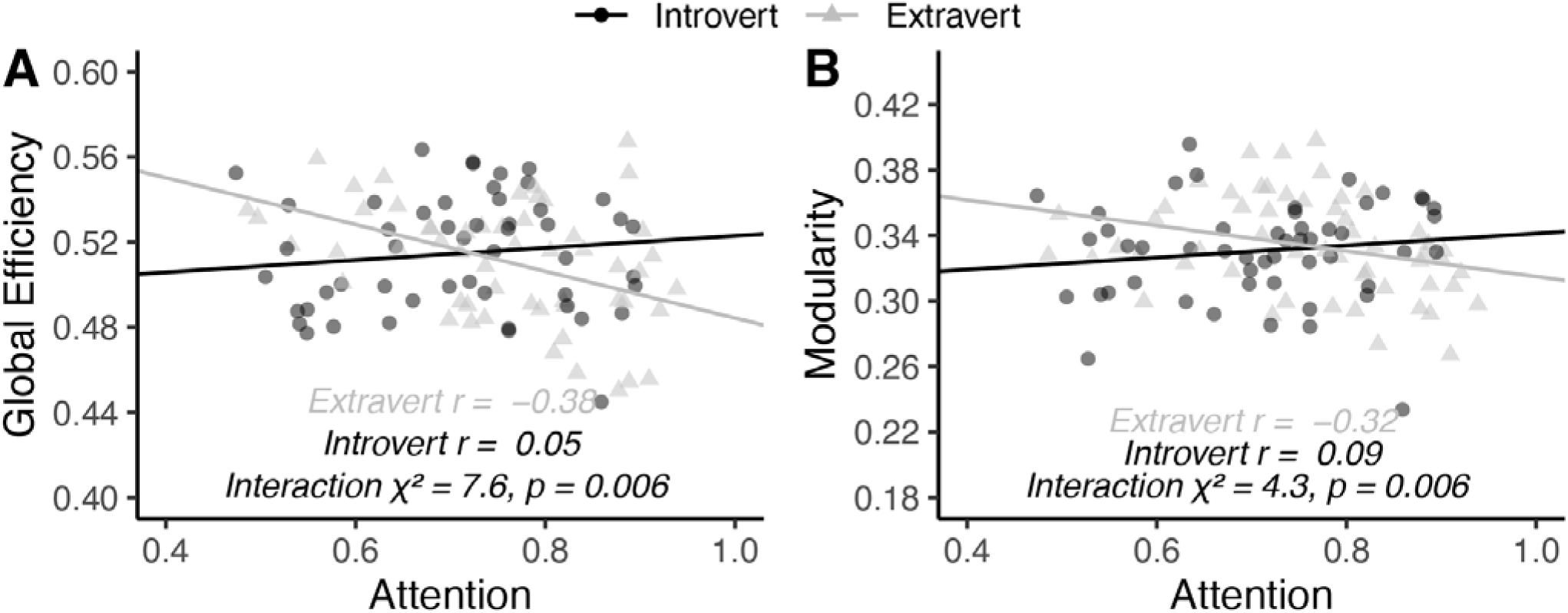
Diverging graph metrics of dynamic network states during focus-on-breath task between introverts and extraverts. Each fMRI scan’s mean attention is correlated against the mean global (A) efficiency and (B) modularity of the individual network connectivity matrices. Introverts (black line) and extraverts (grey line) exhibit differing trends as analyzed by linear mixed effect modeling of interaction effects. Separately analyzed Pearson’s r values are included inset. Points indicate individual fMRI scan values of extraverts (black circles) and introverts (grey triangles).

### 3.3. Extraverts’ and Introverts’ Between-Network Connections to Salience Network Differ During Higher Focused Attention

To uncover which of the three networks contributed to the global efficiency and modularity changes in the dynamic functional networks, we assessed the top 20% whole-brain edges from the previous analysis to characterize their distribution within and between the networks (Figure 3). Notably, the number of external (or, between-network) edges of the SN was found to differ between introverts and extraverts. We observed that increasing mean attention score yielded a greater number of external SN edges during high focus compared to introverts (χ*^2^* = 3.98, *p* = 0.046, Figure 3F). There did not appear to be an extraversion effect on the number of internal edges of the triple networks – the CEN, DMN, and SN (Figure 3A,B,C); but the diverging interaction effect was most pronounced in how the three networks connected to external nodes, with the salience network being the only significantly different network between introverts and extraverts (Figure 3D,E,F).

**Figure 3.**
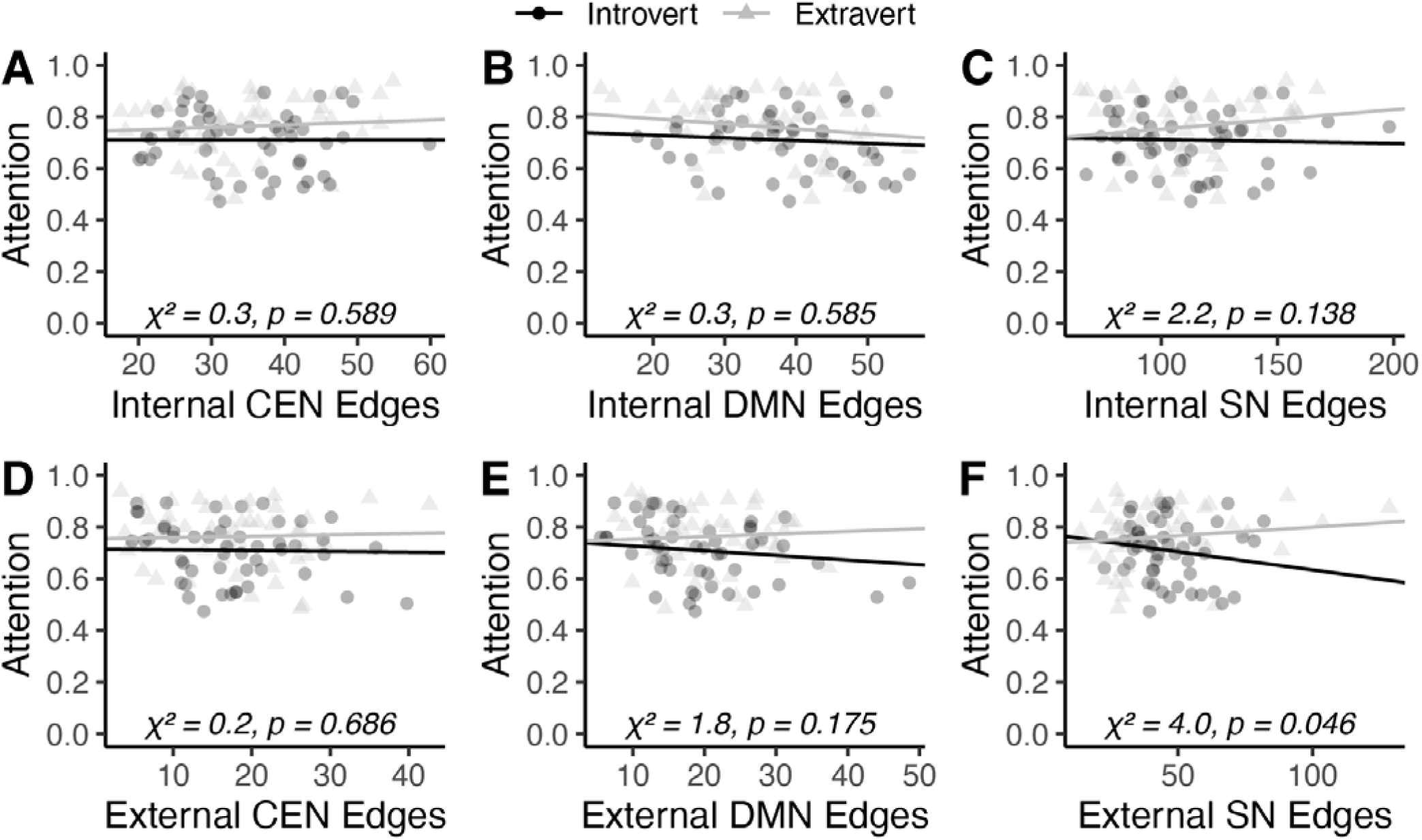
Extraverts and Introverts differ during focus-on-breath meditation in between-network connections of the salience network. Linear mixed effect analyses of the average attention per scan was predicted by an interaction between extraversion and the strength of the top 20% thresholded connections. (A) – (C) show the number of intra-network (internal) edges within the CEN, DMN, and SN. (D) – (F) show the number of inter-network (external) edges from the CEN, DMN, and SN. Each solid black and grey line represent the resulting fixed effect lines for extraverts and introverts respectively and the interaction effect χ^2^ and p-value is displayed inset. Each black circle and grey triangle represents one (of three total scans) for individual extraverts and introverts respectively. Abbreviations: central executive network (CEN), default mode network (DMN), salience network (SN).

### 3.4. Extraverts and Introverts Exhibit Opposing Connectivity Strengths to Achieve Higher Focused Attention

In addition to characterizing the number of edges associated with the triple networks (Figure 3), we also assessed the strengths of the top 20% whole-brain edges. Our results show an interaction between extraverts and introverts for both internal network connections (CEN: χ*^2^* = 6.9, *p* = 0.009, Figure 4A; DMN: χ*^2^* = 3.8, *p* = 0.050, Figure 4B; SN: χ*^2^* = 4.5, *p* = 0.033, Figure 4C) and external network connections (CEN: χ*^2^* = 5.8, *p* = 0.016, Figure 4D; DMN: χ*^2^* = 6.5, *p* = 0.011, Figure 4E; SN: χ*^2^* = 5.1, *p* = 0.023, Figure 4F). Specifically, extraverts exhibit far stronger connectivity strengths of the triple networks – CEN, DMN, and SN – in both internal (Figure 4A,B,C) and external (Figure 4D,E,F) connections associated with higher focus-on-breath. Conversely, introverts tended to exhibit negative connectivity strengths for both internal and external edges of the triple networks – CEN, DMN, and SN. Thus, extraverts’ connectivity strengths increase while introverts’ connectivity strength decrease when participants report optimal attentional focus.

**Figure 4.**
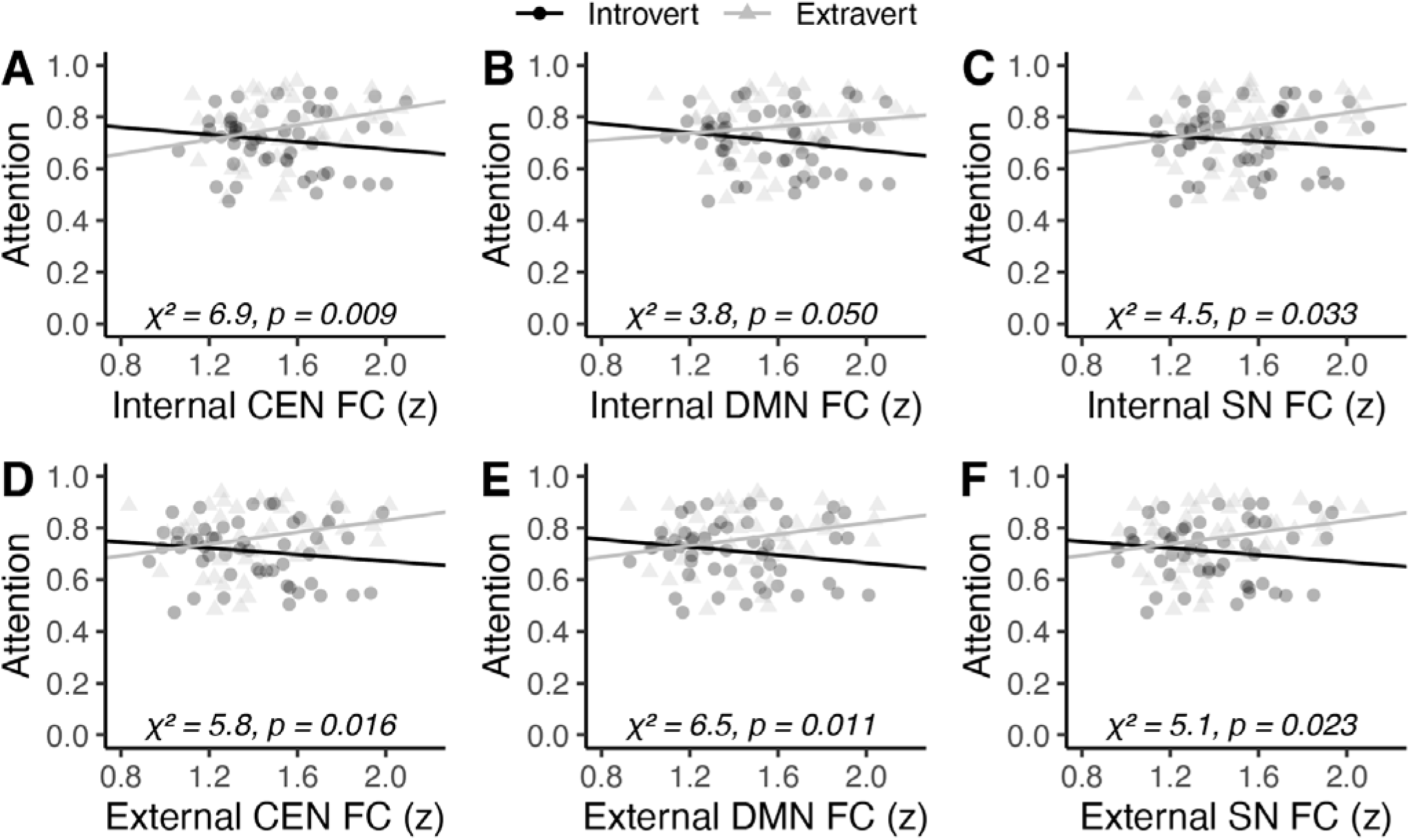
Extraverts exhibit stronger top 20% z-transformed edges in internal and external connections of the CEN, DMN, and SN during high focused attention. Linear mixed effect analyses of the average attention per scan was predicted by an interaction between extraversion and the strength of the top 20% thresholded connections. (A) – (C) show the strength of intra-network (internal) edges within the CEN, DMN, and SN. (D) – (F) show the strength of inter-network (external) edges from the CEN, DMN, and SN. Each solid red and blue line represent the resulting fixed effect lines for extraverts and introverts respectively and the interaction effect χ and p-value is displayed inset. Each black circle and grey triangle represents one (of three total scans) for individual extraverts and introverts respectively. Abbreviations: central executive network (CEN), default mode network (DMN), salience network (SN).

## 4. Discussion

As hypothesized, introverts were more negatively affected by distracting music during meditation, which was most pronounced when the music was rated as having high arousal (Figure 1). Dynamic functional connectivity analysis found that extraverts’ connectomes were less globally efficient and less modular during higher focused attention especially in the presence of distraction (Figure 2). To uncover how the three key networks (CEN, DMN, and SN) differed between extraverts and introverts to support focused-on-breath attention, the number of top 20% connections (Figure 3) and the strength of these connections (Figure 4) showed differences between extraverts and introverts. Specifically, extraverts tended to show higher numbers of external salience network connections (Figure 3F) and greater connectivity strengths of the three networks in the context of higher reported focus-on-breath (Figure 4). Interestingly, introverts exhibited the opposite relationship, such that high focus-on-breath was associated with fewer external salience network connections and decreased connectivity strengths of the triple networks. This divergent relationship between attentional focus, functional connectivity, and broader network structure differences ultimately support Eysenck’s theory of personality that suggests extraverts and introverts have low and high cortical arousal, respectively (Eysenck, 1963).

Extraverts demonstrated superior inhibitory control compared to introverts. Introverts exhibited worsened ability to focus-on-breath with increasing distraction, such that performance was best in the absence of distraction (i.e., no music), was worse during low arousal music, and worst during high arousal music (Figure 1). Importantly, extraverts were able to maintain high attentional focus during high arousal music. This is in line with prior studies which found that extraverts exhibit greater inhibitory control than introverts (Hahn et al., 2015; Prabhakaran et al., 2011). These results also support past literature which assessed task performance in the presence of background music where extraverts performed better than introverts during high arousal music (Cassidy & MacDonald, 2007). However, contrary to other research indicating differential performance in the absence of distraction (Mistry, 2015), we did not observe differences in attentional focus between extraverts and introverts during the no music condition (Figure 1). As such, our results – along with past literature – suggest differences exist between extraverts and introverts in inhibitory control ability in the presence of distraction.

Extraverts’ brain networks are less modular and less globally efficient when they perform inhibitory control, which likely serves to protect against distraction and maintain focused attention on breath (Figure 2). Prior research found that trials of the Stroop task associated with longer reaction times and more errors were associated with brain networks greater in global efficiency and transitivity (an index of the amount of clustering present in the network) (Spielberg et al., 2015). Our results seemingly concur with this prior finding because if our paradigm’s high focus was equated to Stroop task trials with faster reaction times, one would expect less modular and less efficient networks (which we observe in Figure 2). Conversely, if we equated our paradigm’s low focus to Stroop task trials with longer reaction times, we would expect high efficiency and higher clustering – which was reported (Spielberg et al., 2015).

Given that extraverts exhibited better inhibitory control which was related to less global efficiency and modularity (our measures of cortical arousal), we propose that trait extraversion – and its underlying brain network connections – may afford extraverts enhanced inhibitory control, either through a natural predisposition or a greater cognitive alignment with such control processes. Extraverts are thought to have low cortical arousal (Eysenck, 1963) and low cortical arousal in rsfMRI is associated with reduced network modularity (Lee et al., 2022). As such, returning to states of low network modularity and low cortical arousal amidst inhibitory control conditions may be more natural to extraverts.

To further investigate how these brain networks may support extraverts’ focused attention on breath, we observe both the number (Figure 3) and strength (Figure 4) of three prominent large-scale brain networks: the central executive network (CEN), default mode network (DMN), and salience network (SN). By reducing modularity and global efficiency, this would suggest that the effects of distracting external stimuli would not be able to permeate throughout the entire brain as more nodes must be traversed prior to disrupting the networks utilized by the task: CEN, SN, and DMN (Figure 5). We surmise that this organization may protect core nodes related to focused attention, allowing requisite brain regions to perform the directed tasks more proficiently. In other words, extraverts appear to isolate task-based from distraction-based processes by capitalizing on inefficient connectivity. Conversely, our results suggest that extraverts’ connectomes may be more modular and more globally efficient during lower attention (i.e., more mind wandering). Indeed, a past study using resting-state fMRI (i.e., possibly similar to a low-focus state) found that high extraversion was associated with more clustered configurations of brain networks that tended towards small-world networks (Gao et al., 2013). Introverts, on the other hand, show virtually null correlations in how efficiency and modularity measures differ by focus (Figure 2) – indicating some other mechanism by which introverts may protect against distraction. Taken together, our results suggest that extraverts connectomes have lower efficiency and modularity which may reduce the ability for external stimuli to promulgate throughout the network.

**Figure 5.**
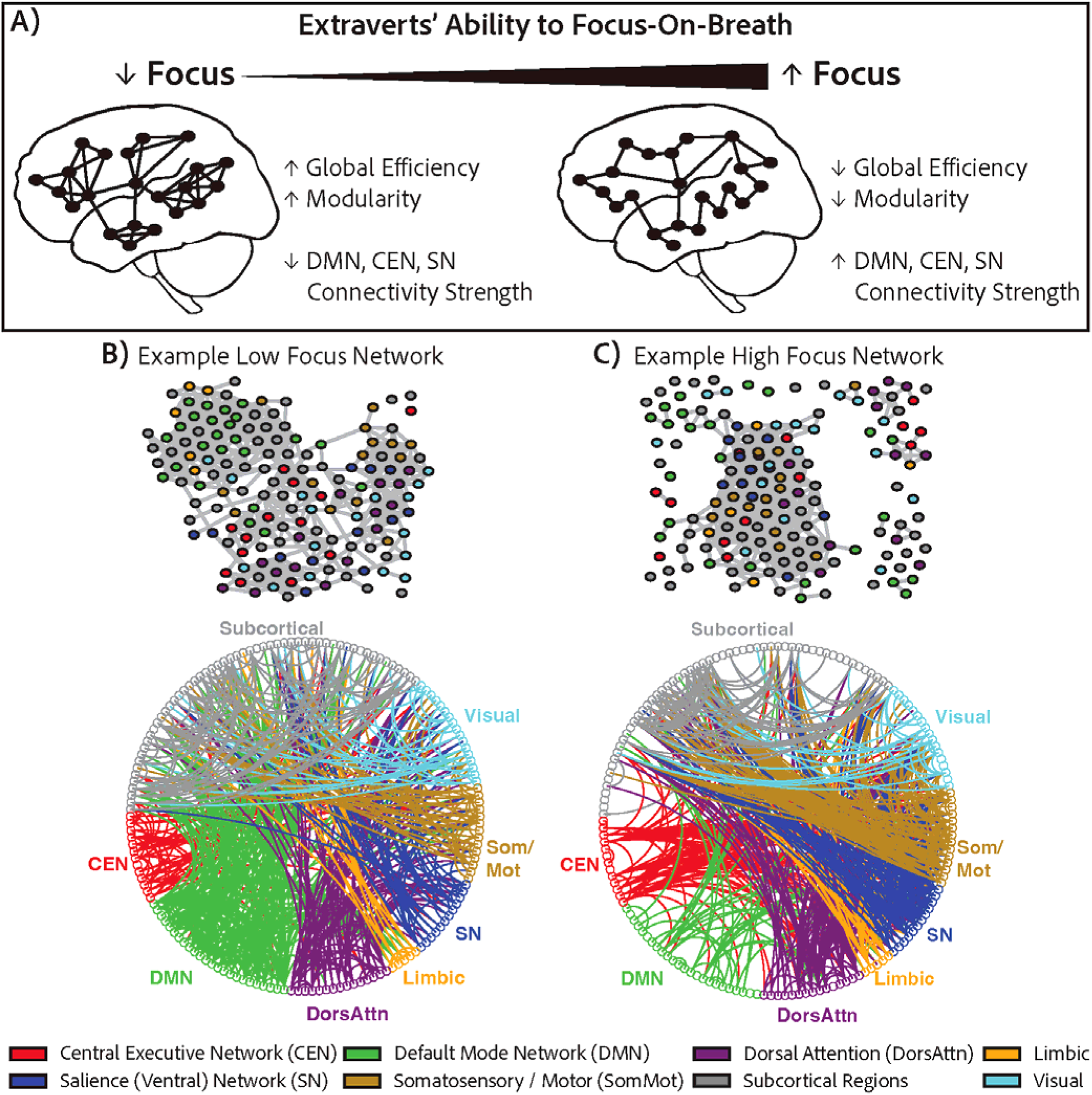
Suggested mechanism by which extraverts protect against distraction. (A) Increased attentional focus is associated with lower global efficiency and lower modularity, such that distracting signals are less able to propagate throughout the network. (B) shows an example of a network with high modularity and global efficiency where the participant exhibited lower focus whereas (C) shows an example of a network with low modularity and global efficiency during high focus. Both (B) and (C) are derived from the same extraverted participant but from high distraction (B) and no music (C) scans. In (B) and (C): the circles denote regions of interest and lines denote an edge of the top 5% (for graphical reasons, rather than top 20%) connectivity strengths with each node color-coded by the network it belongs to according to the Schaefer atlas.

Extraverts form a greater number of salience network connections (Figure 3F) and stronger connections across the CEN, DMN, and SN (Figure 4). The salience network is typically related to orienting or directing one’s attention. For example, the insula, a major salience network hub, is involved in interoception of internal processes – such as focusing on one’s breath (Wang et al., 2019). As the salience network activates to direct one’s attention to target interoceptive sensations (e.g., breath), this competes with the predominant internal mind-wandering processes of the DMN (Ganesan et al., 2022). In line with the hypothesis that extraverts’ have lowered cortical arousal (Eysenck, 1963), the salience network must increase the number of external connections to support higher focused attention. Simultaneously, the extravert’s network reconfigures to be less efficient and less modular such that distracting stimuli (e.g., music) and the elicited processes that occur at subsequent sensory nodes are less able to engage with or shake the predominance of the salience network’s functional connections. Furthermore, extraverts also exhibited stronger intra- and inter-network connectivity measures of the CEN, DMN, and SN to facilitate high focus-on-breath (Figure 4). Alongside extraverts’ less efficient and less modular networks amidst distraction, these stronger connectivity measures may represent an adaptive defense against distraction – protecting key functional nodes critical to performing cognitive tasks (e.g., the central executive networks) from undesired connections and by preventing the breaking of current connections. This may help explain past research that observed strengthened DMN, CEN, and SN inter-network connectivity in resting-state functional connectivity post-meditation sessions (Bremer et al., 2022), which may be an indication of enhanced attentional focus.

This divergent relationship between attentional focus and functional connectivity supports Eysenck’s (1963) theory of personality that suggests extraverts and introverts have low and high cortical arousal, respectively. This divergence between extraverts’ and introverts’ triple networks’ connectivity suggest the possibility for different cognitive processes to achieve focused attention meditation amidst music distraction. The DMN is known to be engaged during introspective thought such as mind wandering (Scheibner et al., 2017) and the decrease in both DMN activity and within-DMN connectivity is linked, in real-time, to focus-on-breath meditation (Brewer et al., 2011; K. A. Garrison et al., 2013). The DMN deactivation is thought to reflect subjective experiences such as “undistracted awareness”, “effortless doing”, or “contentment” (K. Garrison et al., 2013). As introverts may, by nature have higher cortical arousal (Eysenck, 1963) and that the contents of introverts’ mind-wandering is typically more self-referential (Davis & Johnson, 1983), DMN deactivation may be prioritized to allow for introverts to achieve focus-on-breath meditation. Furthermore, the higher cortical arousal of introverts may also mean external musical distraction is particularly exigent such that decreasing triple network connectivity strengths prevents distracting audio stimuli on propagating through the network. As such, decreased DMN connectivity may be the primary mechanism by which introverts achieve focus-on-breath meditation amidst distraction.

The SN and CEN (a.k.a. fronto-parietal network) are integral to focus-on-breath meditation – which may be how extraverts achieve focused attention meditation amidst distraction. The SN directs one’s attention to interoceptive processes (e.g., breath) (Ganesan et al., 2022) and the CEN is involved in cognitive control processes, such as attention and working memory (Menon & D’Esposito, 2022). Both the SN and CEN increase in functional connectivity to support focus-on-breath meditation (Guu et al., 2023; Hasenkamp & Barsalou, 2012). Extraverts’ SN and CEN networks exhibit increased functional connectivity strength (Figure 4), as well as broadened external SN connectivity (Figure 3). Given that the contents of extraverts’ mind-wandering is typically more extrinsic (Davis & Johnson, 1983), activating SN and CEN resources may be met with less resistance as they increase their functional connectivity. Furthermore, rather than reducing connectivity strength to support focus-on-breath meditation, extraverts may instead modify their connectomes to be less efficient and less modular to dynamically protect against distraction (Figure 2). As such, increased SN and CEN connectivity may be the primary mechanism by which extraverts achieve focus-on-breath meditation.

Previous research has shown that hyper- or hypo-functional connectivity has been associated with various neurological and psychiatric disorders. While introversion-extraversion is a personality trait and not a disorder, there are some interesting parallels that may hint at how these personality traits may manifest. For example, autism spectrum disorder is often associated with introversion (Schriber et al., 2014; Vuijk et al., 2018) and children on the spectrum show hyper-functional connectivity (Li et al., 2020). Conversely, attention-deficit/hyperactivity disorder has been associated with both extraversion (Gomez & Corr, 2014; Stanton & Watson, 2016) and hypo-functional connectivity (Castellanos & Aoki, 2016). Our findings help corroborate these relationships to indicate that extraversion and introversion may be dissociated by baseline functional connectivity, where optimal attentional performance may be achieved by altering the magnitude of connectivity in opposing ways depending on the personality type. Furthermore, this supports the notion that a meaningful relationship exists between the extraversion personality dimension, attentional control, and functional network dynamics.

Our results suggest that extraverts may have some cognitive affinity to focused attention meditation. It appears that extraverts may be especially suited for focused attention meditation – particularly with how their functional connectomes reorganize and strengthen to protect against distraction. Indeed, past research has found that extraverts are more likely to try mindfulness meditation (van den Hurk et al., 2011), have more positive attitudes towards meditation (Ryakhovskaya et al., 2025), and have more meditation experience (Ryakhovskaya et al., 2025). While our present study suggests a level of adeptness for extraverts in performing focused attention meditation, another study did not find that extraversion was predictive of individual preference for focused attention meditation (Tang & Braver, 2020).

Although additional research will be needed to confirm these findings and extend them to other task contexts, the idea that personality type differentially engages brain networks may complicate analyses and interpretation of results for neuroimaging studies. However, this may also help explain the high heterogeneity (Aguirre et al., 1998) and replication problems (Kelly & Hoptman, 2022) common to neuroimaging experiments. Future neuroimaging research may benefit from characterizing personality type as a means to understand individual differences in cognition and its associated neural networks.

Overall, the results demonstrated that extraverts exhibit higher focus-on-breath despite the presence of background music, whereas introverts were negatively impacted. This reduced inhibitory control in introverts was hypothesized to occur due to high levels of cortical arousal that make them sensitive to environmental stimuli (Eysenck, 1963). Here, we provided evidence in support of this hypothesis, as introverts reduced functional connectivity to achieve optimal attentional focus. Conversely, it was hypothesized extraverts have low levels of cortical arousal (Eysenck, 1963), and they were observed to increase functional connectivity to achieve optimal attentional focus.

These results not only support Eysenck’s theory, but it extends personality theory to associate extraversion with attentional control and functional connectome dynamics. To explain why extraverts were not negatively impacted by distraction, extraverts were able to dynamically reorganize their connectomes to produce less globally efficient and less modular networks. This indicates that distracting stimuli would travel through more nodes before reaching other important (attention based) nodes, which proffer protection against distraction. Therefore, extraverts increased “cortical arousal” as indexed by both the strength and number of functional connections to isolate attention from distracting stimuli. Conversely, introverts deployed a different method of attentional control, in which they reduced “cortical arousal” as indexed by lower connectivity strength and a reduced number of connections to weaken the effects of distracting stimuli on attentional control. However, introverts did not exhibit any significant change in network topography (global efficiency or modularity) that would help lessen (or isolate) the negative impact of distraction. As such, introverts were distracted by music during meditation.

## Acknowledgements

Thanks to Dr. Huaqing Jin for his valuable insight in the data analyses of this project. We would also like to thank the Neuroscape Network for their generous support of this research.

## Notes

### Competing Interest Statement

The authors have declared no competing interest.

