## SupplementalTable1 for "High extraversion enhances attentional control through dynamic network reorganization"

### Supplement

Table S1 – Extraversion-Introversion questions of the Big Five Personality Traits Questionnaire. Participants rate these statements below on a five-point Likert scale. * denotes questions that should be reverse-scored (i.e., 1 = 5, 2 = 4).

| I see myself as some who… | Strongly Disagree (1) | Disagree (2) | Neutral (3) | Agree (4) | Strongly Agree (5) |
| --- | --- | --- | --- | --- | --- |
| Is talkative |  |  |  |  |  |
| Is reserved* |  |  |  |  |  |
| Is full of energy |  |  |  |  |  |
| Generates a lot of enthusiasm |  |  |  |  |  |
| Tends to be quiet* |  |  |  |  |  |
| Has an assertive personality |  |  |  |  |  |
| Is sometimes shy, inhibited * |  |  |  |  |  |
| Is outgoing, sociable |  |  |  |  |  |
